# Sensory Processing and Categorization in Cortical and Deep Neural Networks

**DOI:** 10.1101/647222

**Authors:** Dimitris A. Pinotsis, Markus Siegel, Earl K. Miller

## Abstract

Many recent advances in artificial intelligence (AI) are rooted in visual neuroscience. However, ideas from more complicated paradigms like decision-making are less used. Although automated decision-making systems are ubiquitous (driverless cars, pilot support systems, medical diagnosis algorithms etc.), achieving human-level performance in decision making tasks is still a challenge. At the same time, these tasks that are hard for AI are easy for humans. Thus, understanding human brain dynamics during these decision-making tasks and modeling them using deep neural networks could improve AI performance. Here we modelled some of the complex neural interactions during a sensorimotor decision making task. We investigated how brain dynamics flexibly represented and distinguished between sensory processing and categorization in two sensory domains: motion direction and color. We used two different approaches for understanding neural representations. We compared brain responses to 1) the geometry of a sensory or category domain (domain selectivity) and 2) predictions from deep neural networks (computation selectivity). Both approaches gave us similar results. This confirmed the validity of our analyses. Using the first approach, we found that neural representations changed depending on context. We then trained deep recurrent neural networks to perform the same tasks as the animals. Using the second approach, we found that computations in different brain areas also changed flexibly depending on context. Color computations appeared to rely more on sensory processing, while motion computations more on abstract categories. Overall, our results shed light to the biological basis of categorization and differences in selectivity and computations in different brain areas. They also suggest a way for studying sensory and categorical representations in the brain: compare brain responses to both a behavioral model and a deep neural network and test if they give similar results.

## Introduction

Neuroscience research has heavily impacted upon recent developments in artificial intelligence (AI) (Hassabis et al., 2017). For example, deep neural networks exploit principles from the visual system in mammals (LeCun et al., 2015) and reinforcement learning, a central approach in modern AI research (Levine et al., 2016; Mnih et al., 2013), originated from theories of animal learning (Pavlov, 1927; Skinner, 1990). However, ideas from more complicated paradigms like perceptual decision making tasks are less used. A large part of recent AI work focuses on automated decision making systems. These are used to help humans take difficult decisions or they take decisions on their own: driverless cars, pilot support systems, medical diagnosis algorithms etc. (Davenport and Harris, 2005; Karanasiou and Pinotsis, 2017; Vatansever et al., 2017). Decisions are often taken in dynamic and challenging environments, under timing constraints, stress or considerable cognitive load (Risko and Gilbert, 2016), thus designing automated decision making systems is challenging too (Grace et al., 2017; Lu et al., 2018). At the same time, humans can effortlessly accomplish many decision making tasks that are deemed difficult for AI systems, e.g. they can flexibly switch between task rules, which AI cannot do. Thus, understanding complex decision making dynamics in the brain and modelling those using deep neural networks could help tackle difficulties faced by artificial systems. To this end, we reanalysed local field potential (LFP) responses recorded during a sensorimotor decision making task using multivariate methods (Representational Similarity Analysis, RSA; Kriegeskorte et al., 2008) and deep recurrent neural networks (RNNs; Hochreiter and Schmidhuber, 1997). LFPs reflect the mass action of millions of neurons within a few hundred microns of each recording electrode. Stimuli were presented across two sensory domains at the same time: motion direction and color.

Monkeys were trained to group different low level sensory features together to form categories based on two rules, categorize random, moving colored dot patterns by direction of motion or by color (Fig 1). Dots moving at 60 and at 120 degrees (on a flat screen) were to be categorized as *upwards* while dots moving at 240 and 300 degrees were categorized as *downwards* (Figure 1). Likewise, they learned to group different blends of red and green colors into *red* vs *green*. They were randomly cued on each trial whether to categorize by motion or color.

**Figure 1.**
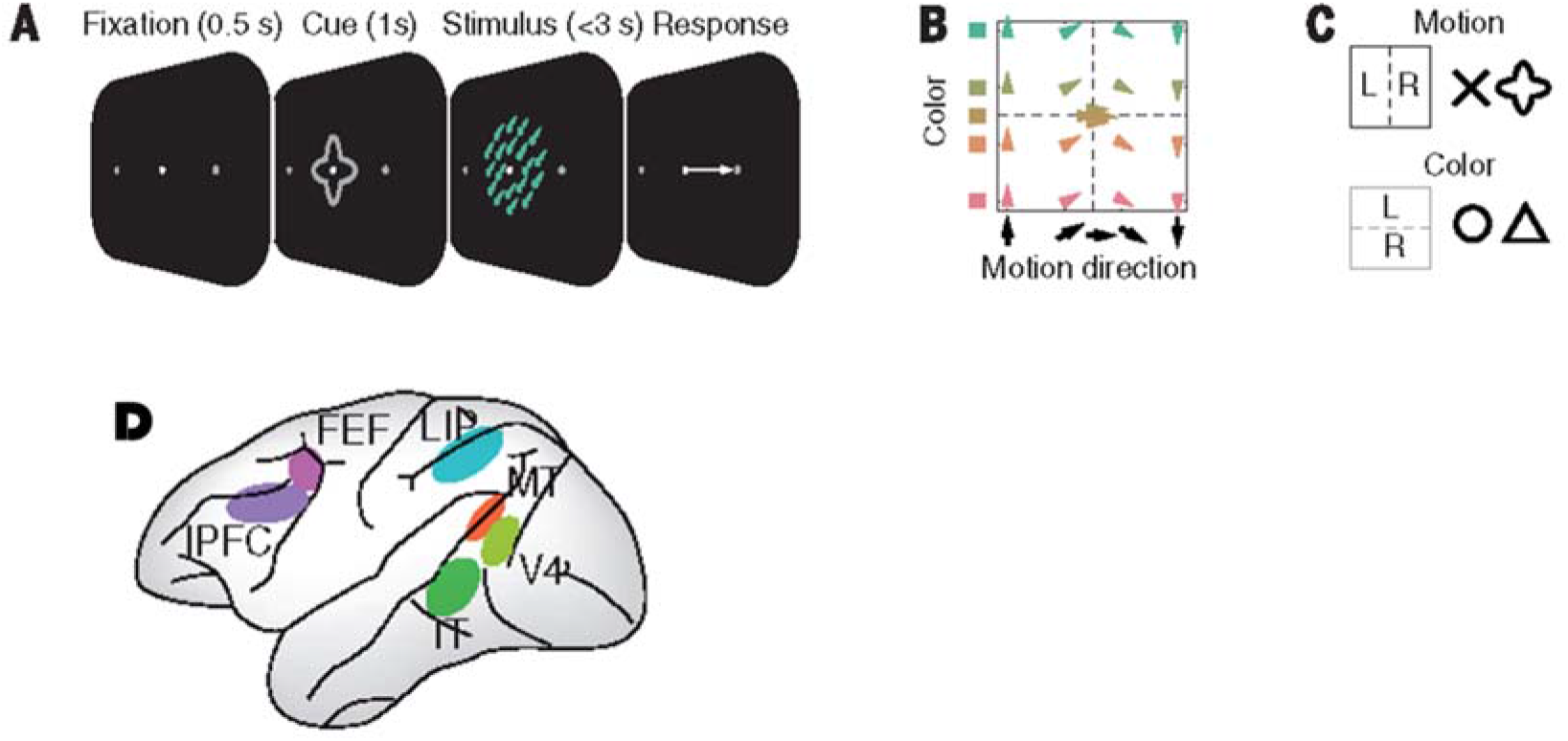
(A) Monkeys categorized the motion direction, or color, of centrally presented, colored random dot stimuli. Before stimulus onset, a central cue indicated which feature to categorize. Monkeys indicated their choice with a leftward or rightward saccade and held central fixation throughout each trial until their response. Monkeys were required to respond within 3 seconds after the stimulus onset. For each trial, we analyzed the data from the stimulus onset to the average response latency (1s to 1.270s) (B) Stimuli systematically covered motion, direction, and color space between opposite motion directions (up and down) and opposite colors (red and green). All stimuli were 100% coherent, iso-speed, iso-luminant, and iso-saturated. (C) Two different cue shapes cued each task. (D) Schematic display of the recorded brain regions. See also (Siegel et al., 2015) for more details.

We investigated how sensory processing and categories were represented by brain dynamics in different cortical areas (Antzoulatos and Miller, 2014; Roy et al., 2014; Wutz et al., 2018) and how these dynamics compare with dynamics in deep RNNs trained to perform the same task. Both kinds of dynamics are important: understanding sensory processing can shed light into the mechanisms of perception. Categorization on the other hand, can help us understand the emergence of rules and abstract thoughts. *Sensory processing* is the result of brain dynamics driven by feedforward sensory input that represent low level features (not categories); here, the exact motion direction and color of the dot patterns, e.g. green dots moving at 60 degrees. Similar dynamics are thought to occur in e.g. deep convolutional networks that perform sensory perception tasks like object classification (Cadieu et al., 2014; LeCun et al., 2015). *Categories* on the other hand, are *functional groupings* of feedforward sensory inputs and thus may depend on distinct dynamics in hierarchies of brain areas, e.g. dots moving at 30 and 60 degrees are categorized as upwards. Categories are thought to result from a combination of feedforward inputs and feedback signals that incorporate prior acquired knowledge (e.g. about task rules, category boundaries, context etc.). In a deep network, it is not clear how categories might be represented. Here, we considered their representations by recurrent activity in deep RNNs in accord with ideas from binding theory (Milner, 1974; Von Der Malsburg, 1994).

To understand neural representations in different brain areas during this flexible decision task, we used two approaches. We computed 1) the similarity of neural representation in a brain area with the geometry of the sensory or category domain represented (we call this *domain selectivity*); 2) The similarity of neural computation performed by a brain area with computations performed by 2 deep RNNs: one trained to distinguish categories (like the behavioural task) and the other to process visual information (we call this *computation selectivity*). The assumption here was that to perform the behavioural task both kinds of computations should take place in different brain areas, i.e. categorization required also sensory processing.

The two approaches we used are distinct. Being selective to a sensory domain (domain selectivity) is not the same as performing computations like sensory processing and abstract categorization (computation selectivity). Domain selectivity refers to representation content only, while computation selectivity characterizes how these representations are manipulated and compared to each other to find their similarities and differences. Also, sensory processing requires integrating sensory inputs while abstract categorization requires combining these integrated inputs with prior knowledge about learned categories. All these computations take time. Thus, understanding which computations each area performs requires analyzing temporal information in brain dynamics. Although distinct, domain and computation selectivity should give similar results. We found this below (see *Results*).

We used RSA to quantify the selectivity of each brain area to different sensory and category information. We found that representations in different brain areas change flexibly depending on whether they contained sensory or category information and whether the monkeys were instructed to categorize color vs motion. Thus, selectivity of different brain areas depended not only on the stimulus represented but also on the task (color vs motion categorization). We then used deep RNNs and LFPs to study the computations that each brain area performed. By comparing network predictions with brain dynamics using RSA and the Bayesian Information Criterion (BIC), we found task-dependent computations in different brain areas. All in all, our work sheds light on the nature of representations and computations during a complex sensorimotor decision making task. This can be useful for building deep neural networks that can take informed decisions in dynamic, real life scenarios. The design of such deep neural networks would extend the analyses presented here. Their layers would be designed to perform computations similar to those performed by cortical networks doing the same task—like the layers in the deep neural networks we considered below.

## Methods

### Subjects and recordings

Experiments were performed in two rhesus monkeys (one male, one female). All procedures followed the guidelines of the Massachusetts Institute of Technology Committee on Animal Care and the National Institutes of Health. Each monkey was implanted with a titanium head bolt to immobilize the head. Following the behavioral training, three titanium recording chambers were stereotactically implanted over frontal, parietal, and occipitotemporal cortices in the left hemisphere. Through these chambers, we simultaneously implanted Epoxy-coated tungsten electrodes in the lateral prefrontal cortex, frontal eye fields, lateral intraparietal cortex, inferotemporal cortex (TEO), visual area V4, and the middle temporal area (MT). Recordings were performed acutely—electrodes were inserted and removed each session. Offline, we extracted the continuous local field potentials (LFPs) by first removing DC offset and line noise, parsing into trials by low-pass filtering at 250 Hz (2nd-order zero-phase forward-reverse Butterworth filter), and resampling at 1 kHz (Van Kerkoerle et al., 2014).

### Behavioral Task

Monkeys were trained on a flexible visuomotor decision making task (Figure 1). All stimuli were displayed on a color calibrated CRT monitor at 100 Hz vertical refresh rate. An infrared based eye-tracking system continuously monitored eye position at 240 Hz. Behavioral control of the behavioral task was handled by the Monkeylogic program (www.monkeylogic.net). The trial began with fixating a fixation target at the center of a screen (500ms; Figure 1A). Fixation was required within 1.2° of visual angle of the fixation point. Then, monkeys were presented with a visual task cue for 1 sec. Time zero corresponded to when the cue appeared. They were four such cues (Figure 1C) - two of them cued the monkeys to categorize the direction of motion (motion task) and the other two cued them to categorize the color of the moving, colored dots (color task). Two different cues were used to signify each task so that we could dissociate information about the appearance of the cues from the tasks they cued. Using two cues for each task allowed us to dissociate neuronal information about the cue (the visual shape of the cue) and neuronal information about the task at hand (motion vs. color).

After the cue, a random dot stimulus was presented centrally on the fixation spot. Stimuli were colored moving random dot patterns with 100% motion coherence (stimulus diameter: 3.2°; dot diameter: 0.08°; number of dots: 400; dot speed: 1.67°/s or 10°/s for half of the recording sessions, respectively). To prevent learning of patterns, new stimuli were generated for each session. There were 7 possible stimulus motion directions and 7 possible colors. 4 possible motion directions spanned the range between −90 and 90 degrees in 60 degree steps (−90º, −30º, 30º,90º) (Figure 1B). In addition, 3 motion directions were placed on (0º) and near (−5º, 5º) the category boundary. Similarly, 4 colors spanned the range between red (90º) and green (−90º) through yellow (0º) in 60º steps (−90º, −30º, 30º, 90º). In addition, three colors were placed on (0º) and near (−5º, 5º) the category boundary. Depending on the cue, the animals categorized either the color (red vs. green) or motion direction (up vs. down) of the stimulus and reported their percept with a left or right saccade. Thus, there were in total (4×4+5) x2 =42 different stimulus conditions. Animals could respond any time up to 3 sec after stimulus onset (Figure 1A). We only considered trials were the animals gave the correct response in our analyses. Animals performance was satisfactory and similar in both tasks. Animals were always rewarded for ambiguous trials with stimuli on the category boundary.

To optimize perceptual homogeneity, all colors were defined in the CIE L*a*b* space and had the same luminance and saturation. The stimulus-response mapping for each task was fixed (Figure 1C). Two saccade targets were displayed 6º to the left and right of the fixation spot throughout the trial. Animals had to respond with one direct saccade to one of these targets.

Note that we here analysed data only from the interval from stimulus onset to the average response latency (1 to 1.270s). In (Siegel et al., 2015), we found that this is the interval that contains motion and color information (just after the stimulus appears and before the choice response). We used this interval to compute LFPs as we were specifically interested in how dynamics in different brain areas represent this information. Understanding its nature is important for developing automated decision making systems, which motivated this paper^1^. We computed LFPs from up to 108 electrodes simultaneously implanted in six cortical areas acutely each day (Figure 1D): MT, LIP, V4, IT, FEF and PFC. For more details about the experiment, see (Siegel et al., 2015).

### Representation Dissimilarity Matrices (RDMs)

Representation Dissimilarity Matrices (DMs) are used to summarize how stimulus information is mapped to brain responses. They capture differences and similarities in brain responses corresponding to different stimuli and provide a characterization of representation content in each brain area. We constructed them following (Kriegeskorte et al., 2008 and Figure 2, upper panel). For each time point (1ms) in the interval between stimulus onset and average response latency (1 to 1.270sec), we averaged LFP time series across all trials corresponding to each of the 42 experimental conditions. Thus we obtained 42 motion and color patterns (over electrodes). We computed the dissimilarity between them (i.e. 1-Pearson correlation).We considered time correlations that are thought to underlie categories according to binding theory (von der Marlsburg, 1994). Alternatively, we could had summarized electrode activity using PCA or considered space correlations. We also averaged over time—and did not consider possible changes of category selectivity over time, e.g. (Kadipasaoglu et al., 2016; Scholl et al., 2014). We thus obtained RDMs of dimension 42×42. This has the advantage that it normalizes for both the mean level of activity and its variability as discussed in (Kriegeskorte et al., 2008). The more similar two sets of LFPs were, the lower the dissimilarity between them was.

**Figure 2.**
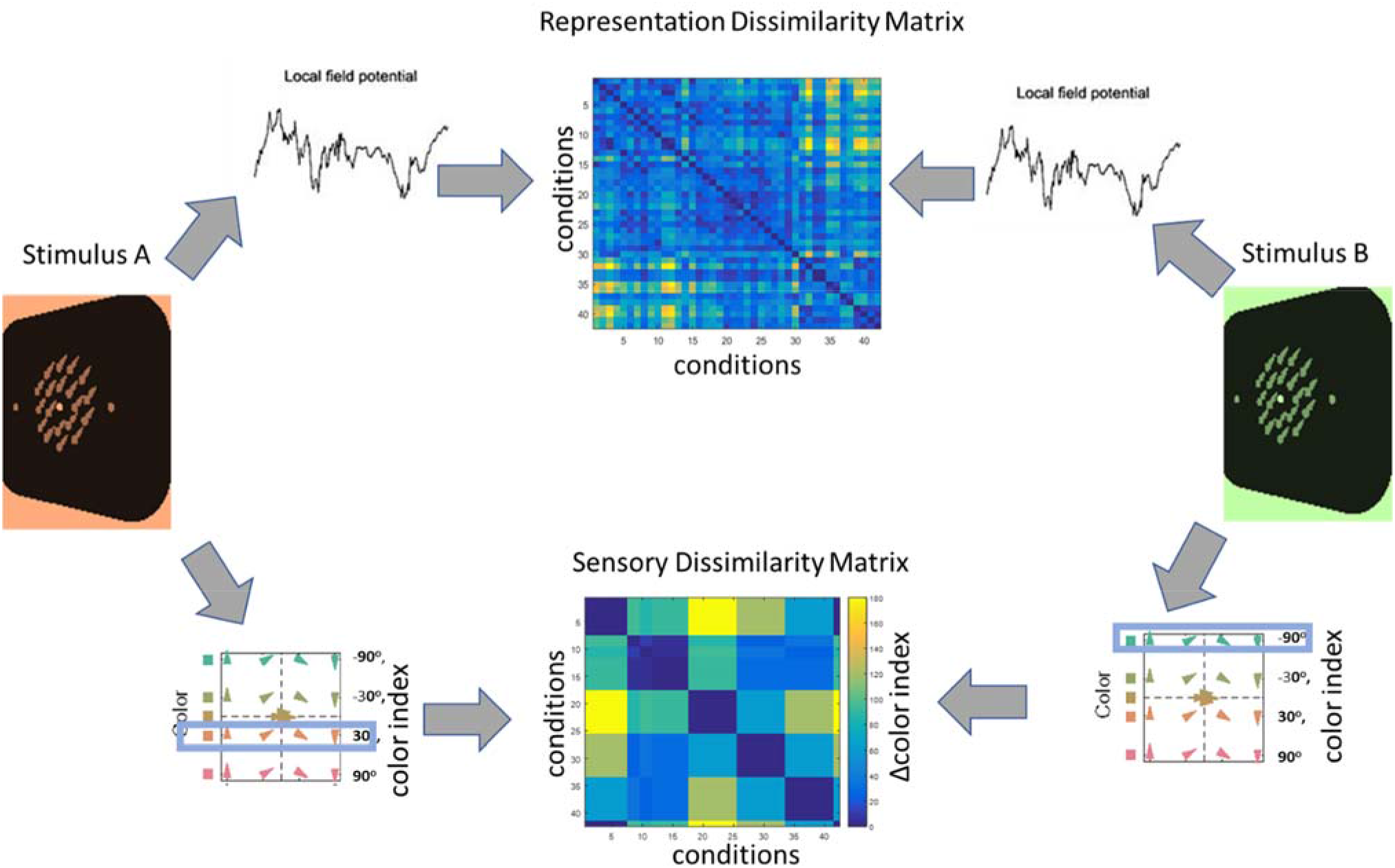
Construction of Dissimilarity Matrices. We grouped LFPs recordings corresponding to conditions (e.g. Stimulus A in the motion task and Stimulus B in the color task, see left and right panels respectively) and computed the correlation distance between them. We repeated the same process for all stimuli and computed pairwise correlations. We then obtained a RDM for each brain area (upper panel) after averaging over time. Constructing a color Sensory Dissimilarity Matrix (color SDM) was similar (lower panel). We associated each condition with an index. We then calculated pairwise index differences for all conditions. Here, the color index was 1 for greenish and 0 for pinkish stimuli.

RDMs corresponding to higher visual areas known to play a role in object recognition, V4 and IT, showed a more pronounced clustering over motion color or motion conditions compared to other areas (structuring by quadrants in V4 and IT RDMs, cf. Figure 3A). This could be explained by their color selectivity (Heywood et al., 1992; Zeki et al., 1991) and stronger retinotopy (Fize et al., 2003; Tootell et al., 1998). To distinguish RDMs obtained above using LFP time series from other kinds of DMs we consider below, we call these matrices *brain* RDMs.

**Figure 3.**
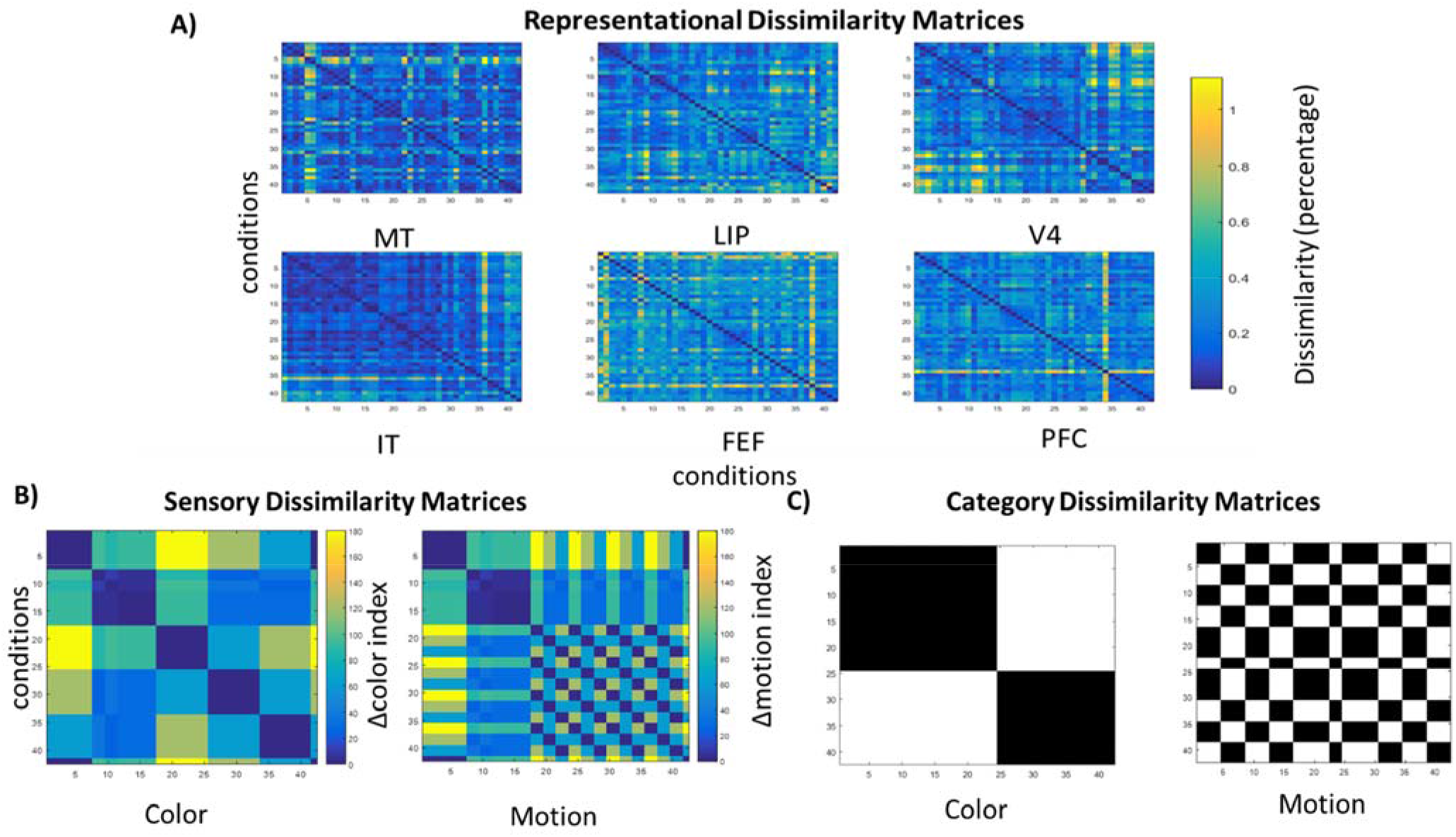
Dissimilarity Matrices (DMs). (A) Representation DMs for the six brain areas we recorded LFPs from. These matrices describe stimulus representations in each area. (B) Sensory DMs in color and motion direction domains. These matrices show pairwise differences between different stimulus values (color and motion indices) and how they are distributed in the respective domains. Differences in color and motion indices are smooth and shown with a continuous jet color map. (C) Category DMs in color and motion direction domains. These matrices show pairwise differences between different categories and the corresponding binary category distribution. Differences in the color and motion categories are shown with a binary color map. All in all, DMs capture the geometry of sensory stimulus and category domains.

### Network RDMs and Sensory and Category DMs (SDMs/CDMs)

Besides brain RDMs, we also constructed RDMs using predictions from deep RNNs trained with LFPs from the same time interval as the one we used for brain RDMs above (1 to 1.270s). These deep RNNs are described in detail below We call the RDMs obtained using RNN predictions, *network* RDMs. We also constructed two other kinds of DMs, that is, sensory and category DMs (SDMs and CDMs(Kriegeskorte, 2011). These matrices illustrate the geometry of the sensory and category domains respectively, i.e., how different stimuli are distributed in the stimulus space and the categories and are sometimes referred to as behavioural models. Also, SDMs are often considered as continuous models, as opposed to categorical models embodied by CDMs. We constructed the color SDM as follows (the process for the motion SDM was similar). Each of the 7 color conditions was associated with a color index. The difference between color indices was calculated between pairs of conditions. These differences formed the color SDM shown in Figure 2 (lower panel). For the color CDM, we only had two conditions (i.e. categories). Here, the color index was 1 for greenish and 0 for pinkish stimuli, cf. Figure 1B.

### Representational Similarity Analysis (RSA)

RSA was introduced by (Kriegeskorte et al., 2008) to assess the similarity between neural activity and predictions from behavioral or computational models. This uses DMs of the sort considered above. Here, we used RSA for two sets of analyses. First, to compare brain RDMs with SDMs and CDMs. Second, to compare brain and network RDMs. The comparison between brain RDMs and SDMs/CDMs allowed us to assess the selectivity of each brain area to different low level visual features (motion direction, color) and their categories. On the other hand, the comparison between brain and network RDMs allowed us to understand the exact computation performed by each brain area in the motion direction and color tasks.

To understand what kind of representations (motion direction vs color, sensory processing vs categorization) were encoded in each brain area, we computed the dissimilarity between brain RDMs and SDMs or CDMs. As in (Kriegeskorte et al., 2008) the dissimilarity between these dissimilarity matrices, known as *deviation*, was the correlation distance (1-Spearman correlation; Spearman was used as it does not require a linear correspondence between these matrices contrary to Pearson correlation, see (Kriegeskorte et al., 2008)). Deviations between RDMs and SDMs or CDMs quantify matches between representation content of brain responses and the geometry of sensory stimulus and category domains (Kriegeskorte, 2011). They measure the correlation distance^2^ between each RDM (describing pairwise differences in patterns of neural activity corresponding to different experimental conditions) and each SDM or CDM (describing pairwise differences in low sensory features or categories corresponding to the same conditions).

In other words, deviations quantify *second-order differences* (i.e. differences of differences): How different are the corresponding pairwise differences in neural activity on the one hand and sensory information or categories on the other. Since SDMs and CDMs describe the geometry of sensory stimulus and category domains respectively, *the smaller the second-order differences, the higher the match between patterns of neural activity and stimulus or category domains*. This, in turn, indicates that the corresponding brain area is more selective (sensitive) to sensory information or categories (i.e., smaller deviations imply more selectivity to the corresponding sensory or category domains).

Each brain RDM was correlated with 1) the SDMs/CDMs and 2) network RDMs. This process yielded between-DM deviations shown in 1) Figure 4 and 2) Figures 4 and 5 in *Results*. Error bars in these Figures denote the standard errors. To estimate these errors bars, i.e. the variability of deviations (had we chosen different stimuli from the same population), we used bootstrap resampling (N=100 repetitions with replacement^3^) and obtained a distribution of correlation values. To test whether two DMs were related, we used fixed effects category-index randomization test. We simulated the null distribution. This has the advantage that it does not require normality and independence assumptions like the Bonferroni correction. 10,000 relabelings were obtained by reordering rows of the DMs. We thus obtained a distribution of correlations with smallest possible significance level *p=10^-4^* (two-tailed probability level relative to the null distribution that the two DMs were unrelated).We assumed a a false positive rate of 5. If the actual correlation we had obtained fall within the top 5% of the simulated null distribution, then we reject the null hypothesis. Rejecting the null hypothesis, that is, finding that RDMs and SDMs/CDMs are related meant that that the corresponding brain area was *selective* to the sensory domain or the category corresponding to the SDM or CDM respectively; while finding that brain and network RDMs were related meant that the brain area performed a similar *computation* to what the corresponding deep recurrent neural network did (sensory processing or categorization). To report inferential statistics we followed (Kriegeskorte et al., 2008). All deviations were significant at the *p<0.05* level except for those which are shown to be non-significant at this level (denoted by “n.s.” at the bottom of the corresponding bars in Figures 4, 5 and 6).

**Figure 4.**
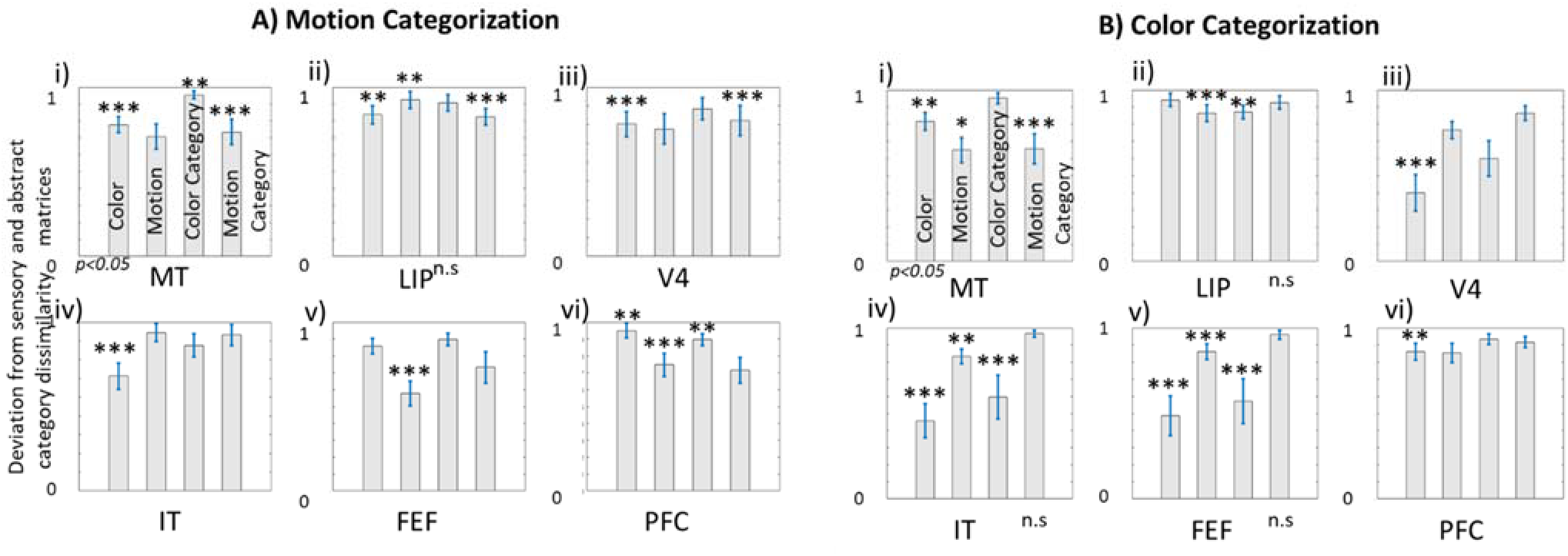
Deviations between RDMs and SDMs/CDMs. (A) Motion categorization task. Each panel depicts deviations between RDM of a brain area and the SDM (“color”, “motion” 1^st^ and 2^nd^ bars from left) or CDM (“color category”, “motion category”, 3^rd^ and 4^th^ bars) respectively. Error bars denote standard errors. All deviations were significant at the p<0.05 level with the exception of those with “n.s” at the bottom(not significant; fixed-effects category-index randomization test, see Methods and (Kriegeskorte et al. 2008)). (B) Same results for the color categorization task. Note that deviation is based on correlation distance, thus smaller bars indicate better similarity between RDMs and SDMs/CDMs. Asterisks above each bar denote the significance level of the corresponding partial correlations.

**Figure 5.**
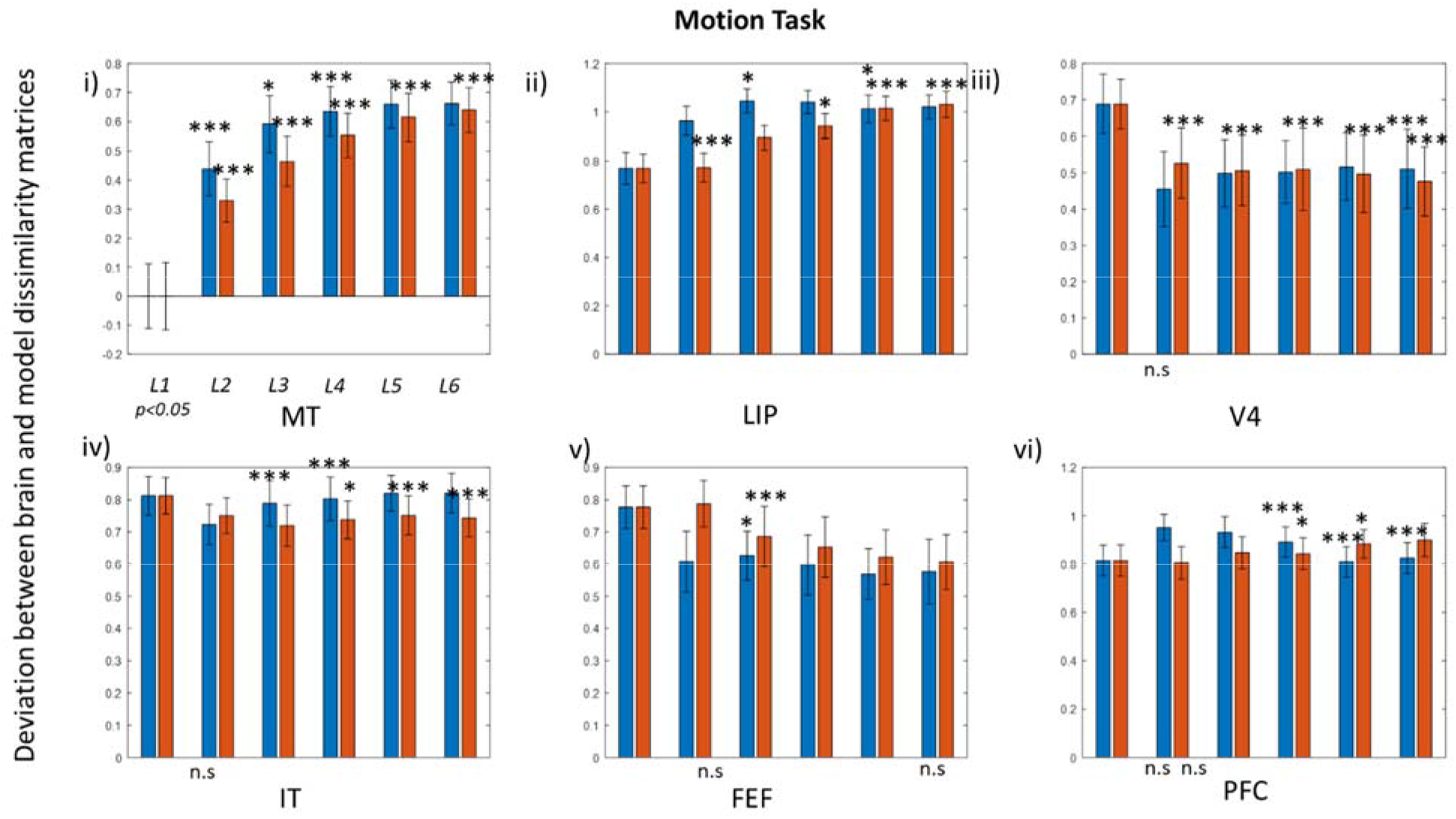
Deviations between brain and network RDMs. Bars in each panel depict deviations between RDM of a brain area and each layer in a deep RNN performing motion processing and categorization. There are six pairs of bars, equal to the number of layers. The left bar in each pair corresponds to deep RNN predictions when the network performs sensory processing, while the right bar corresponds to predictions during categorization. Error bars denote standard errors. All deviations were significant at the p<0.05 level with the exception of those with “n.s” at the bottom (not significant; fixed-effects category-index randomization test, see Methods and (Kriegeskorte et al. 2008)). Note that deviation is based on correlation distance, thus smaller bars indicate better similarity between brain and network RDMs. Asterisks above each bar denote the significance level of the corresponding partial correlations.

**Figure 6.**
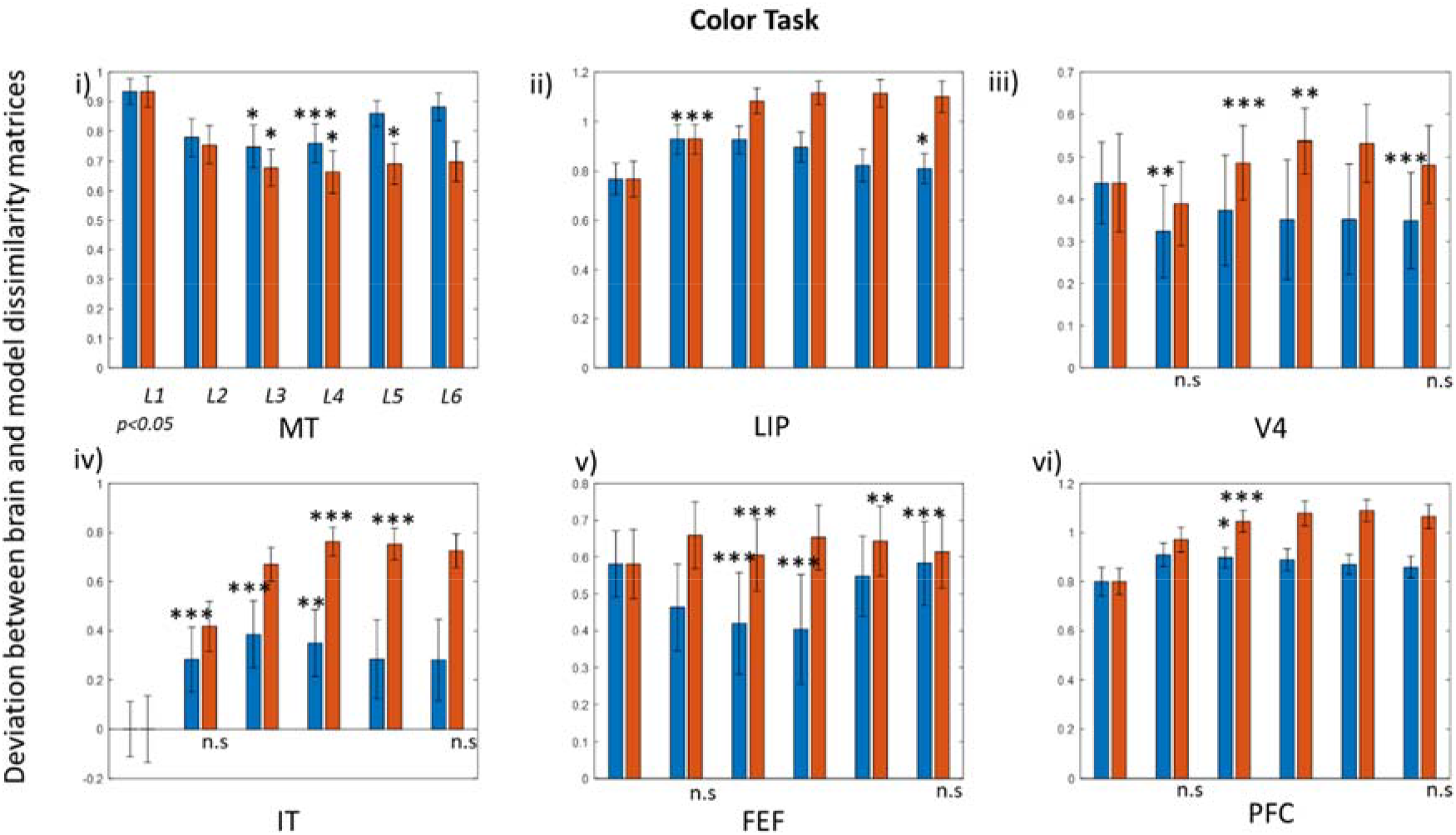
Deviations between brain and network RDMs for color processing and categorization. This is similar to Figure 5 where the deep RNN has learned to process and categorize color as opposed to motion direction stimuli.

### Deep recurrent neural networks (RNNs) for sensory processing and categorization

We here used a popular class of deep RNNs called Long-Short Term Memory (LSTM) networks. LSTMs have several advantages, for example, they prevent catastrophic effects due to vanishing and exploding gradients (Hochreiter and Schmidhuber, 1997). We used LSTMs as they are known to be able to learn temporal correlations of the sort present in a categorization task like the one considered here, very efficiently (Pearlmutter, 1989). Testing LSTMs against other deep architectures (e.g. CNNs) was beyond the scope of this work as we were interested in recurrent dynamics which are modeled by LSTMs. This dynamics is important for categorization as, according to binding theory (Milner, 1974; Von Der Malsburg, 1994), category representations are thought to rely on temporal correlations.

We used LSTMs as proxies of large cortical networks to shed light on the computations various brain areas performed and trained them using LFP recordings and the same sensory and category labels we used for constructing color and motion SDMs and CDMs (see above).

All RNN implementations were done in Keras using Tensorflow backend and RMSprop optimization. The LFPs used for training were obtained from the same time interval as the one used for brain RDMs (1 to 1.270sec). All RNN networks had six LSTM layers equal to the number of brain areas from which we recorded and a last dense layer (with a softmax nonlinearity) that could predict output conditions (classification). The number of units in each layer were (38,28,28,38, 38,38) respectively. The number of units and other model parameters were found using a pretraining approach that is described in detail in *Supplementary Results*. For regularization, we used dropout in the first and last LSTM layers with 40% rate. This randomly omits a fraction of connections between two network layers during training. To optimize the learning rate, we also constrained the size of network weights with max-norm regularization (max weight= 5), see also *Supplementary Figure 1*.

We ensured all RNNs achieved satisfactory performance: they performed better than linear regression (see *Supplementary Results* and *Supplementary Figures 1 and 2*). To prevent overfitting, we stopped training while train and testing accuracy where increasing. In all cases, we stopped training when accuracy was well above chance, see *Supplementary Figure 4*.

## Results

We examined sensory processing and categorization in cortical activity and in deep RNNs. Our goal was to understand the role of different brain areas involved and how brain dynamics changed when task demands changed. We reanalysed data from a visuomotor decision making task performed by two monkeys (Siegel et al., 2015). They categorized the motion direction and color of moving, colored, random dot stimuli (Figure 1). We analysed local field potentials (LFPs) during the interval between stimulus onset and the animal’s response (average reaction time, RT, i.e. 1 to 1.270 sec, *Methods*). We chose this interval because it contains motion and color information (Siegel et al., 2015).

### Distributed representations of sensory domains and categories in the frontoparietal brain network

To understand brain dynamics during the flexible decision task, we first analysed the *selectivity* of different brain areas to low level visual features (actual motion direction, color) and to motion and color categories. We chose RSA to compute brain area selectivity because it allows us to evaluate the similarity of the same brain responses to both the geometry of stimulus spaces and deep neural network predictions in the same way. We can thus distinguish between domain selectivity (selectivity of a brain area to a sensory or category domain) from computation selectivity (selectivity to either sensory processing or categorization) that requires manipulation of neural representations (see Table 1 and *Discussion*). We first constructed representation dissimilarity matrices (RDMs) for each brain area, which we call brain RDMs (see *Methods* and Figure 3A).

**Table 1.**
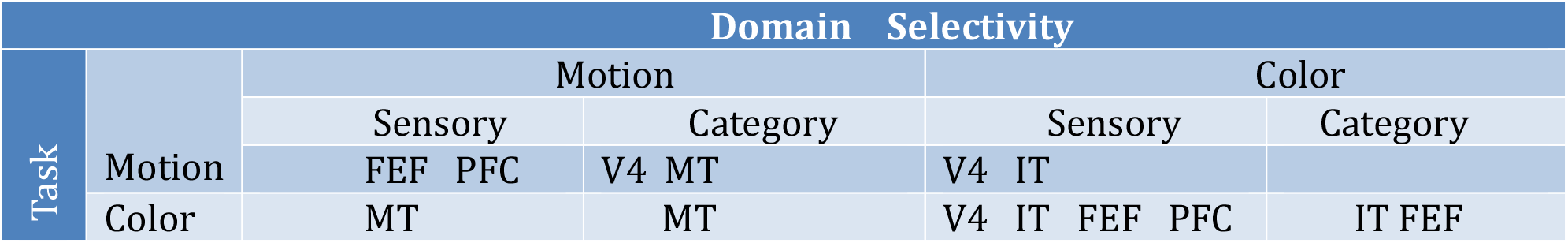
Domain selectivity of each cortical area depending on task.

To understand what kind of representations (motion direction vs color, sensory processing vs categorization) were encoded in each brain area, we computed the deviation, ie. the dissimilarity between brain RDMs and SDMs/CDMs. Since both motion direction and color information were simultaneously present, it is not clear which sensory feature was represented in each brain area. Brain RDMs were computed based on neural activity, therefore representations of sensory features co-existed with representations of categories.

To test the hypothesis that a brain area was selective to a certain sensory or category domain, we tested whether a particular deviation was significant or not (*Methods*). A significant deviation implies that the corresponding RDM and a behavioral model, that is, SDM/CDM matrices are correlated. This means that pairwise differences in neural activity are correlated with pairwise differences in stimulus or category domains. Thus, patterns of neural activity induced by sensory processing or categorization reflect the geometry of these domains. Significance was assessed using a fixed effects category-index randomization test and post hoc analyses using multilinear regression (*Methods*).

We asked whether there was an effect of task on representation, i.e., if representations in different brain areas depended on whether animals were cued to categorize motion vs color. Thus, before computing the deviations, we split each of the dissimilarity matrices of Figure 3 into submatrices corresponding to trials where the monkey was cued to categorize motion or color respectively. The results of our analysis are shown in Figure 4 for the motion (Figure 4A) and color (Figure 4B) categorization tasks respectively.

In all these figures (panels of Figure 4), bars denote the deviation between RDMs (neural activity) and SDMs/ CDMs (stimuli or behavioral model). Each of the four bars in each panel shows the deviation of the RDM of a brain area from the color and motion SDMs (first and second bars in plots of Figure 4, see also Figure 3B) and color and motion CDMs (third and fourth bars in plots of Figure 4, see Figure 3C). Error bars denote standard errors. RDMs were correlated with SDMs/CDMs (significant deviations at the *p<0.05* level) with the exception of LIP RDM and color CDM in the motion categorization task (Figure 4Aii) and LIP, IT and FEF RDMs and motion CDM in the color categorization task (Figures 4Bii, 4Biv and 4Bv). To contrast the predictive power of each SDM/CDM separately over and above all other SDMs/CDMs, we used multilinear regression.

The significance of the corresponding partial correlations is denoted by asterisks over the bars in Figure 4. This allowed us to assess the significance of each deviation separately. We asked which among the candidate SDMs and CDMs matched better a particular brain RDM. We thus found the sensory or category domain for which a certain brain area was more selective. This was the domain that had the smallest deviation, which remained statistically significant after controlling for variance shared with the other SDMs and CDMs.

Considered together, the results of Figures 4A and 4B show that selectivity of most brain areas depended on the task. That is, selectivity changed depending on the rule the monkey followed in each trial (categorize color vs motion). This is because for most brain areas the bars (deviations) that were the smallest were different (in the corresponding panels in 4A and 4B). Consider e.g., the second and fourth bars in 4Aiii vs the first and third bars in 4Biii. These are the smallest deviations between V4 activity and SDMs/CDMs. Since the second and fourth bars in 4Aiii were the smallest this means that V4 was more selective for motion (2^nd^ bar) and motion categories (4^th^ bar). Similarly because the first and third bars in 4Biii were the smallest this means that V4 was more selective for color (1^st^ bar) and color categories (3^rd^ bar). Thus V4 selectivity changed in the two tasks. This was also the case for most brain areas. The only exceptions were MT and IT which showed the same selectivity regardless of task. Our results are summarized in Table 1 and are discussed below in detail…”

Based on deviations, V4 showed more motion and motion category selectivity during the motion task (the deviations between its RDM and the motion and motion category DMs are smaller than the other two deviations; second and fourth bars in Figure 4Aiii). Similarly, V4 showed more color and color category selectivity during the color task (first and third bars in Figure 4Biii are smaller than the other two). Multilinear regression revealed that in the motion task, deviations with motion CDM (*w=2×10^-3^, p<0.001*) and color SDM (*w=2×10^-4^, p<0.001*) remained significant after controlling for the other regressors. In the color task, the deviation with the color SDM (*w=4×10^-4^, p<0.001*) remained significant. Considering together which deviations were the smallest in each task and significant, we found that V4 preferred color in both tasks and motion category in the motion task. Thus, V4 selectivity changed with the task.

FEF and PFC selectivity also changed with the task. FEF preferred motion and motion category in the motion task (second and fourth bars in Figure 4Av are smaller than the other two) and color and color category in the color task (first and third bars in Figure 4Bv are smaller than the other two)^4^. Also deviations between FEF RDM and motion CDM were not significant in the color task. (see “n.s.” below the rightmost bar in Figure 4Bv). This means that FEF was not selective (insensitive) for motion categories in the color task. After controlling for other regressors, deviation with motion SDM remained significant in the motion task (*w=4×10^-4^, p<0.001*). Also all deviations (except the n.s. with motion CDM) remained significant in the color task using multilinear regression (color CDM: *w=2×10^-2^, p<0.001*, color SDM: *w=4×10^-4^, p<0.001*, motion SDM: *w=2×10^-4^, p<0.001*). Considering which deviations were the smallest and significant, FEF appeared selective for motion in the motion task and color and color categories in the color task.

PFC was more selective for motion than color during the motion task (second and fourth bars are smaller than the other two in Figure 4Avi). Which domain PFC was more selective for, during the color task was not clear based on deviations (they were similar and above 90%, see Figure 4Bvi). However, using multilinear regression for the color task, the only significant deviation was the one with color SDM (*w=2×10^-4^, p<0.01*). Also, this deviation remained significant in the motion task (*w=4×10^-4^, p<0.01*) along with deviations with motion SDM (*w=10^-3^, p<0.001*) and color CDM (*w=5×10^-2^, p<0.01*). Thus, it appeared that PFC was more selective for motion in the motion task and color in the color task.

Based on deviation values, it is not clear what the LIP selectivity was (deviations are similar and above 90%, see Figures 4Aii and 4Bii). Interestingly, LIP wasn’t selective to the categories that were irrelevant for the task. Only the relevant categories e.g. motion categories in the motion task etc., showed significant deviations, see “n.s.” below the deviation from the color CDM in the motion task; third bar from the left in Figure 4Aii and similarly “n.s.” below the deviation from the motion CDM in the color task; rightmost bar in 4Bii. Also, controlling for the rest of the regressors, we found that in the motion task, deviations with the motion SDM (*w=-4×10^-4^, p<0.01*), color SDM (*w=4×10^-2^, p<0.01*) and motion CDM (*w=7×10^-2^, p<0.001*) were significant. In the color task, deviations with color CDM (*w=4×10^-2^, p<0.01*) and motion SDM (*w=10^-3^, p<0.001*) were significant.

To sum up, selectivity for most brain areas depended not only on the stimulus but also on the domain of categorization, i.e. the task. V4, FEF and PFC selectivity seemed to follow the task. They preferred the motion or color domains in the motion and color task respectively. However, two areas were the exception to this rule: MT and IT. For them, selectivity depended only on the stimulus presented, and the task was irrelevant. Based on deviations, MT was selective for motion and motion category in both tasks (second and fourth bars from left are smaller than the other two in both Figures 4Ai and 4Bi). Interestingly, the deviation with motion SDM did not remain significant when using multilinear regression (*w=10^-4^, p=0.8*). All other correlations remained significant in the same task (color CDM: *w=-10^-2^, p<0.01*, color SDM: *w=3×10^-2^, p<0.001*, motion CDM: *w=2×10^-2^, p<0.001*). In the color task, most deviations remained significant too (color SDM: *w=10^-4^, p<0.01*, motion SDM: *w=10^-4^, p<0.05*, motion CDM: *w=2×10^-2^, p<0.001*). Taking all above results together, MT was more selective for motion categories in both tasks and motion in the color task (at a lower significance level p<0.05, see the single asterisk in Figure 4Bi).

Similarly, based on deviations, IT was selective for color and color category in both tasks (first and third bars are smaller than the other two in both Figures 4Aiv and 4Biv)^5^. Deviations remained significant only with color SDM in the motion task (*w=4×10^-4^, p<0.001*) and color CDM, color SDM and motion SDM in the color task (color CDM: *w=8×10^-3^, p<0.001*, color SDM: *w=2×10^-3^, p<0.001*, motion SDM: *w=6×10^-5^, p<0.01*). Taking above results together, IT was more selective for color in both tasks and color categories in the color task.

### Differences in sensory processing and categorization by the cortical network revealed with a deep recurrent neural network (RNN)

Above, we found that different brain areas were selective to different sensory and category domains. We compared brain activity to the geometry of the sensory or category domain represented. Below we focused on the differences in the semantic content of representations in different brain areas and asked the following question: what kind of computation did a brain area perform more, sensory processing or abstract categorization? The assumption here was that to perform the behavioral task (categorize) some brain areas in the frontoparietal cortical network implicated in the task, would perform sensory processing, i.e. they would represent sensory information independently of the learned categories.

To answer this question, we used deep RNNs based on LSTMs. Technical details about these RNNs are included in *Methods* and *Supplementary Results*. It should be noted that they were *not* precise descriptions of brain *anatomy*. Instead, they merely provided simulations of brain *computations* (sensory processing or categorization). These simulations resulted in predictions of brain activity that were then compared to brain activity that was measured. After training, their accuracy was 2-3 times above the chance level (*Supplementary Figure 4*).

Training was implemented using supervised learning. LFP responses were provided as input. The output was conditions (labels) corresponding to each experimental condition. After training, a comparison between predictions from separate layers of the deep RNN with brain activity suggested what kind of computation each area performed (see below). We trained 2 variants of the same deep RNN for each task. The first deep RNN performed sensory processing (21 motion direction and 21 color conditions); and the other categorization (2 motion direction and 2 color categories). Below, we will call the first variant *sensory* RNN and the other *category* RNN. We will also discuss how activity from a certain brain area compared to their predictions. If it was more like the sensory RNN, then we concluded that that the brain area was performing sensory processing. Otherwise, categorization.

We used LFP recordings as input from those areas whose selectivity did not depend on task. We found above that most brain areas changed their selectivity depending on task (motion or color). The only exceptions were LFP recordings from MT and IT. Thus, we used LFP recordings from MT and IT as input to the deep RNN. Note that the current paradigm involves flexible decision making in two sensory domains, which is different from sensory perception experiments often considered in the literature where information propagates in a feedforward fashion only and selectivity of different brain areas does not change during the experiment, see e.g. (Cadieu et al., 2014; Yamins et al., 2014, and see also Discussion). Also, MT and IT are well known to be sensitive to motion and color respectively (this is also their selectivity that we found in Figure 4). Thus, we trained the networks with LFP recordings from MT and IT as input for the motion and color tasks respectively. We also used the corresponding sensory and category indices (depicted in SDMs and CDMs, cf. Figure 3) as labels^6^. In *Supplementary Results* we also discuss training the deep RNN performing the color task using MT (the brain area where motion and color information rose first) rather than IT responses as input.

Different network weights were learned while training the deep neural networks with MT and IT recordings as input. These weights depended on the nature of the computation performed by the networks (sensory processing vs categorization) and sensory domain (motion vs color). After training, we performed the same analysis as in Figure 4, where we replaced motion and color SDMs and CDMs with RDMs constructed using deep RNN predictions (*Methods*). These network RDMs used RNN predictions from different layers. We computed the deviations between the RDMs corresponding to each brain area and RNN layers (brain and network RDMs) for each task and sensory domain. The results of our analysis are shown in Figures 5 and 6, for motion and color task respectively. The format of these Figures is similar to Figure 4^7^. They reveal which computation (sensory processing or categorization) each brain area was more selective for.

We assumed that the deep RNN learns the task through forming distributed representations that spread across all layers. In the next paragraph, we will see use RSA and brain and network RDMs to study what kind of computations were performed by each brain area. Each panel in Figures 5 and 6 corresponds to a different brain area (same order as Figure 4). There are six pairs of bars in each panel, equal to the number of deep RNN layers. The left and right bars in each pair in the panels of Figure 5 (in the following “left” and “right” bars for simplicity), depict deviations obtained by training the deep RNN to classify motion direction based on sensory information and categories respectively; while the bars in Figure 6 show the corresponding deviations obtained for the color task. These deviations were obtained after computing the RDMs for each brain area and each network layer and correlating them. Error bars in Figures 5 and 6 were obtained by bootstrap resampling and denote standard errors (see *Methods*).

Deviations between brain and network RDMs quantify matches between representation content of brain responses and network predictions (Kriegeskorte et al., 2008). Similarly to the discussion of Figure 4 above, they also quantify second-order differences, i.e. differences between pairwise differences in neural activity on the one hand and network predictions on the other (for the same pairs of experimental conditions). The smaller the second-order differences, the higher the match between patterns of neural activity and network predictions. This, in turn, implies that the brain area is performing the computation that the deep RNN is performing (sensory processing or categorization) during the two tasks. To sum up, the results of Figures 5 and 6 allow us to conclude which computation each area was more selective for (sensory processing vs categorization).

We used the same approach as in Figure 4. We focused on the smallest deviations between brain RDMs and network RDMs that describe deep RNN predictions for different stimuli and tasks. We asked if they were significant and tested if differences in neural activity were correlated with differences in network predictions. If they were, this means patterns of neural activity and deep RNN predictions were similar. This allowed us to conclude what sort of computation a certain brain area was more selective for. This was the computation performed by the deep RNN whose layers satisfied the following constraints: 1) they had the smallest deviations; 2) these deviations were significant and 3) they remained significant when computing partial correlations.”

To confirm that this RNN predicted brain activity better we also used model comparison. We compared two general linear models (GLMs) with significant layers of either the sensory or category RNN as regressors. We scored them (their predictive power against brain activity) using the difference in the Bayesian Information Criterion, *ΔBICsc*, corresponding to each GLM; this is a standard approach, see (Schwarz, 1978). If *ΔBICsc>0* this confirms that the category RNN predicts better brain activity, while for *ΔBICsc<0*, the sensory RNN is a better predictor. Also, the absolute value of this difference suggests how strong the evidence in favor of the winning model is. If |*ΔBICsc*|>*2*, there is good evidence in favor of the winning model, |*ΔBICsc*|>*6*, suggests strong evidence and |*ΔBICsc*|*>10* is very strong (Kass and Raftery, 1995).

Similarly to Figure 4, partial correlations quantified the significance of deviations for each individual RNN layer^8^ when controlling for variance shared with other layers and the alternative deep RNN. This is denoted by asterisks over the bars in Figures 5 and 6. Also, statistical significance of unpartialed correlations was obtained by randomization tests (see also Methods). In Figures 5 and 6, we report inferential statistics following (Kriegeskorte et al., 2008). Most layers showed deviations with brain activity that were significant at the p<0.0001 level. The only exception was the RDM corresponding to the second layer of the network trained to perform sensory processing that showed significant deviations only with MT, LIP and FEF RDMs in the motion task (second left bars from the left in Figures 5i, 5ii and 5iii). Also, the same RDM was significantly correlated only with MT, LIP, V4 and IT RDMs in the motion categorization (second right bars in Figures 5i, 5ii, 5iii and 5iv). Also, the RDMs corresponding to the second and last layers of the network trained to perform color categorization showed significant deviations only with MT and LIP RDMs (second and last right bars in Figures 6i and 6ii). Below, we discuss the selectivity of each area in detail.

We found that MT was more selective for categorization during both the motion and color tasks. First, in the motion task, MT activity resembled more layers of the RNN, performing categorization compared to the RNN that was trained to perform sensory processing. Deviations with 5 layers of the category RNN remained significant as opposed to 3 layers in the sensory RNN: To denote layers, we use the notation LXC and LXS, where X is the layer number in the deep RNNs performing categorization and sensory processing respectively. Then, multilinear regression yielded, L2C: *w=0.5, p<0.001*, L3C: *w=1.7, p<0.001*, L4C: *w=-2.1, p<0.001*, L5C: *w=2.1, p<0.001*, L6C: *w=2, p<0.001* vs. L2S: *w=0.6, p<0.001*, L3S: *w=-1.3, p<0.05* L4S: *w=1.9, p<0.001*. Also, deviations with the category RNN were smaller and activity was more similar to brain activity: in 4 out of 6 pairs the right bars are smaller; deviations, *D,~37-57%*, second to fifth right bars in Figure 7i; vs *D~42-60%*, second to fifth left bars (the first layer was the input layer hence deviation is zero). Comparing GLMs comprising layers of the category and sensory RNNs as regressors using BIC, we found that the difference in the evidence between them was *ΔBICsc>10*. This confirmed that evidence in favor of the category RNN was very strong.

In the color task, MT activity also resembles more activity in the category RNN (deviations were smaller, right bars were smaller than left bars, *D~70% vs. D~80-90%*, see third to sixth right and left bars in Figure 6i.) 3 of these deviations remained significant after controlling for the rest of the layers (L3C: *w=0.4, p<0.05*, L4C: *w=3.5, p<0.05*, L5C: *w=-21, p<0.05*). Also, deviations with 3 layers of the sensory RNN were significant (L2S: *w=0.6, p<0.001*, L3S: *w=-1.3, p<0.05* L4S: *w=1.9, p<0.001*). Note that these corresponded to higher deviations hence activity in the sensory RNN was not as similar to brain activity as activity in the category RNN. Also, in this case *ΔBICsc>6*, which suggested strong evidence again in favor of the category RNN. To sum up, MT preferred categorization in both tasks. It also seemed to prefer more motion rather than color categorization.^9^

We then considered IT. In the motion task, which computation IT might prefer was not clear because deviations, *D*, were weak and error bars suggest that deviations between the two RNNs can overlap (*D>70*, third to sixth right bars in Figure 5iv). Deviations with 3 layers in the category RNN remained significant when controlling for other regressors, (L4C: *w=0.2, p<0.05*, L5C: *w=-0.2, p<0.001*, L6C: *w=0.3, p<0.001*), while deviations with 2 layers remained significant in the sensory RNN (L3S: *w=0.4, p<0.001*, L4S: *w=-0.3, p<0.001*). When computing the difference in evidence between the 2 alternative RNNs using BIC we found evidence in favor of the sensory RNN, *ΔBICsc>-10*. In the color task, IT preference for sensory processing was found to be strong using deviations and was confirmed using BIC, i.e. *ΔBICsc>-10* too (*D~30%*, second to sixth left bars in Figure 6iv; note that the first layer was the input layer hence deviation is zero). 3 layers in the sensory RNN remained significantly correlated (L2S: *w=0.16, p<0.001*, L3S: *w=-0.03, p<0.001*, L4S: *w=0.023, p<0.01*), while 2 layers remained significant in the category RNN (L4C: *w=-2.5, p<0.001*, L5C: *w=19.3, p<0.001*). Taking all above results together, IT seemed to prefer sensory processing in both tasks.

Based on deviations, V4 showed preference towards sensory processing in the color task and motion categorization in the motion task. At the same time V4 selectivity for ccolor ategorization was also high (all bars in Figure 6iii are small indicating high selectivity for all computations, but sensory RNN bars, on the left of each pair, are smaller). Deviations with the second and last layer of the category RNN were non-significant. When contrasting each individual layer using partial correlations, we found that 2 layers of the sensory RNN (L2S: *w=0.04, p<0.05*, L6S: *w=0.04, p<0.001*) and 2 layers of the category RNN were significantly correlated (L3C: *w=-0.4, p<0.001*, L4C: *w=7.9, p<0.05*). Evaluating the difference in evidence between the alternative RNNs, we found *ΔBICsc>-10*, which is very strong evidence in favor of the sensory RNN. On the other hand, which computation V4 was more selective for during the motion task was not clear based on deviations (they were similar for both the sensory and the category RNN in Figure 5iii). However, after contrasting each individual layer against others using partial correlations we found that only 1 layer of the sensory RNN was significantly correlated (L6S: *w=0.7, p<0.001*) vs. 5 layers of the category RNN (L2C: *w=0.3, p<0.001*, L3C: *w=-0.5, p<0.001*, L4C: *w=-0.2, p<0.001*, L5C: *w=0.4, p<0.001*, L6C: *w=-0.7, p<0.001*). Also, the difference in RNN evidence was in favor of the category RNN, *ΔBICsc>10*.

FEF showed clear preference for sensory processing during both tasks (*D~60%* and *D~45%*, second to sixth left bars in Figures 5v and 6v respectively). This was confirmed by the *ΔBICsc>-10* in both tasks.3 layers of the sensory and 2 layers of the category RNNs were significantly correlated in the color task after controlling for each regressor independently (L3S: *w=-0.18, p<0.001*, L4S: *w=0.3, p<0.001*, L6S: *w=-0.09, p<0.001*; L3C: *w=-1.0, p<0.001*, L5C: *w=15.7, p<0.05*). At the same time, 1 layer in the sensory and the category RNNs remained significantly correlated in the motion task (L3S: *w=0.17, p<0.05*, L3C: *w=-1.7, p<0.001*).

LIP showed weak selectivity (*D>70%*) that changed with the task: it was more selective for categorization during the motion task (*D~85%*, second to fourth right bars in Figure 5ii). 4 layers of the category RNN remained significantly correlated during the motion task after controlling for each individual layer independently vs. 2 layers of the sensory RNN (L2C: *w=0.9, p<0.001*, L4C: *w=0.7, p<0.05*, L5C: *w=-1.0, p<0.001;* L6C: *w=1.8, p<0.001* vs. L3S: *w=-0.7.7, p<0.05*, L5S: *w=-2.1, p<0.05*). *ΔBICsc> 6* which suggested that the category RNN was a better predictor of brain activity. On the other hand, based on deviations, LIP appeared selective for sensory processing during the color task. Selectivity was weak (*D~85%*, third to sixth left bars in Figure 6ii) and only 1 layer of each of the 2 RNNs remained significantly correlated after computing partial correlations (L6S: *w=0.1, p<0.05, vs*. L2C: *w=0.7, p<0.001*). There was evidence in favor of the sensory RNN, *ΔBICsc>-2*. To sum up, LIP seemed to prefer categorization in the motion task, and sensory processing in the color task.

Finally, PFC also showed weak selectivity (*D>70%*) and showed mixed preference. During the motion task, PFC activity resembled activity in layers of both networks (in the third and fourth pairs the right bars are smaller while in the fifth and sixth the left bars, Figure 5vi). After contrasting each individual against others using multilinear regression we found that 3 layers of the sensory RNN remained significantly correlated as opposed to 2 layers in the category RNN (L4S: *w=1.3, p<0.001*, L5S: *w=8.1, p<0.001*, L6S: *w=-8.2, p<0.001* vs. L4C: *w=-0.7, p<0.05*, L5C: *w=-0.6, p<0.05*). Evaluating RNN evidence, we also found support in favor of the sensory RNN, *ΔBICsc >-10*. In the color task, PFC selectivity was also weak. Based om deviations only, PFC appeared to weakly prefer sensory processing more (*D~80%*, second to sixth bars in Figure 6vi). Contrasting each layer against others, we found that only 1 layer from each RNN remained significant (L3S: *w=0.17, p<0.05, vs*. L3C: *w=-1.7, p<0.001*). Then comparison of the 2 alternative RNNs using BIC offered support in favor of the category RNN, *ΔBICsc >6*. All above results are summarized in Table

**Table 2.**
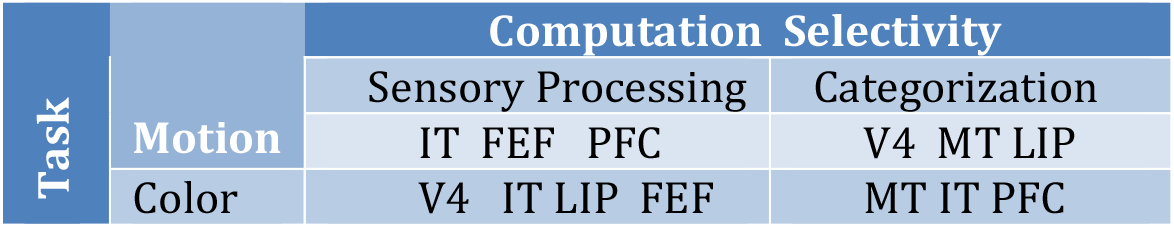
Computation selectivity for each cortical area and task.

In general, for both tasks, we found similar results using the two approaches considered here, that is, comparison of neural activity to 1) the geometry of the sensory or category domains (domain selectivity), and 2) to predictions of deep RNNS, (computation selectivity). V4 showed preference towards sensory processing in the color task and motion categorization in the motion task. Similarly, we had found V4 preference to the color domain in both tasks and motion categories in the motion task. MT was more selective for categorization during both the motion and color tasks. In accord with that, we had found MT was more selective for the motion category domain in both tasks. FEF showed clear preference for sensory processing during both tasks. Similarly, we had found selectivity for motion in the motion task and color in the color task. PFC seemed to prefer more sensory processing in motion task and categorization in the color task. Similarly, we had found selectivity for the motion domain in the motion task and the color domain in the color task. Finally, IT seemed to prefer sensory processing in both tasks which also coincided with its domain selectivity: IT was more selective for color in both tasks and color categories in the color task.

## Discussion

In previous work, we found that complex decision making signals in the brain resulted from an interaction between sensory signals propagating in feedforward paths along the cortical hierarchy with feedback decision signals from frontal and parietal areas (Siegel et al., 2015). We recorded LFPs from a large cortical network comprising six brain areas, V4, IT, MT, LIP, FEF and PFC and found that these recordings contained information about cue, task, motion direction and color categories. In that paper, we also used raw information measures (variance explained) and focused on the temporal dynamics of cue, task, motion, color and choice information. Here, we reanalysed the same LFP data using RSA and deep RNNs. Our goal was to understand brain dynamics underlying complex sensorimotor decisions in more detail. Motivated by automated decision making systems that have no explicit cue or task information, we limited our investigation to motion and color representations. Understanding such representations and how they differ between brain areas is important for developing automated systems that could work in consort with humans, e.g. pilot support systems that would use brain activity corresponding to these representations for rapid decision making.

To understand neural representations in different brain areas, we used two approaches. We computed 1) the similarity of neural representation in a brain area with the geometry of the sensory or category domain represented (which we call *domain selectivity*; motion vs color vs motion categories vs color categories). 2) The similarity of neural computation performed by a brain area with predictions from 2 deep RNNs: one trained to distinguish categories (like the behavioural task) and the other to process visual information (*computation selectivity*). The assumption here was that to perform the behavioural task both kinds of computations should take place in different brain areas, i.e. categorization required also sensory processing.

These approaches are distinct. Being selective to a sensory domain (domain selectivity) is not the same as performing computations like sensory processing and abstract categorization (computation selectivity). Domain selectivity refers to representation content only, while computation selectivity characterizes how these representations are manipulated and compared to each other to find their similarities and differences. Also, sensory processing requires integrating sensory inputs while abstract categorization requires combining these integrated inputs with prior knowledge about learned categories. All these computations take time. Thus, understanding which computations each area performs requires analyzing temporal information in brain dynamics. To study computation selectivity we used neural networks trained on time resolved data. This is another difference between domain and computation selectivity. While domain selectivity was assessed through correlations that did not contain temporal information, computations cannot be understood with such simple, time-independent, statistical techniques.

First, we discuss domain selectivity. To compute this, we used representational similarity analysis (RSA; Nili et al., 2014). RSA is a standard method for comparing brain activity to behavioural and neural models. There are other methods that we could have used for this comparison, like encoding analysis, pattern component modelling (Diedrichsen and Kriegeskorte, 2017), time-frequency mutual information (Baraniuk et al., 2001; Gong et al., 2018) or decoding methods (Christophel et al., 2012; Jazayeri and Movshon, 2006). Using RSA, that is, representational distances between experimental conditions as summary statistics has the advantage that we can assess the similarity of the same brain responses to the geometry of stimulus spaces and deep neural network predictions in a simple, unified way. This, in turn, leads to a detailed understanding of representation content and neural computation. This also provides insights to efficient deep neural network training. Note that knowing the domain selectivity also suggested which brain area to choose as input for neural network training.

Domain selectivity is summarized in Figure 4. Interestingly, for most brain areas it depended not only on the stimulus but also on the domain of categorization. It switched between the two domains depending on task (motion direction or color categorization). This is a surprising result, not previously shown to the best of our knowledge in a flexible decision making task. Also, related work in the literature usually focuses on sensory perception only, and does not normally involve flexible switching between sensory domains, contrary to the paradigm considered here. Indeed, a similar change in domain selectivity was found in studies where attentional state was directed either at location or a specific feature dimension (Bichot et al., 2015; Desimone and Duncan, 1995; Liu et al., 2007). This led to differences in behaviour, improving performance related either to spatial or feature attention respectively. Improved performance is thought to result from changes in either single neuron activity (Reynolds et al., 1999) or population tuning (Serences and Boynton, 2007) and representations (Guggenmos et al., 2015).

The only exception to the above rule were areas MT and IT. Their selectivity depended on stimulus only and not the domain of categorization. MT was more selective for motion categories in both tasks and motion in the color task. Similarly, IT was more selective for color in both tasks and color categories in the color task. Our results are in accord with classical considerations about functional specialisation of these areas which suggest that MT and IT are sensitive to visual motion and color features respectively. Our results also suggest that MT contains direction-selective neurons as it is known from the classical work of (Dubner and Zeki, 1971) and other studies, e.g. (Osborne et al., 2004). They are also consistent with (Brouwer and Heeger, 2009) who could *not* decode color from MT+ activity. Regarding IT, our results confirm the important role of this area in object recognition in general and color processing in particular (DiCarlo et al., 2012; Komatsu et al., 1992).

As said, selectivity of all other areas was flexible and changed when the task or domain of categorization changed. Specifically, we found that FEF, PFC and V4 selectivity followed the task at hand. FEF appeared selective for motion in the motion task and color and color categories in the color task. PFC was more selective for motion in the motion task and color in the color task. V4 preferred color in both tasks and motion category in the motion task. (Xiao et al., 2006) reported evidence suggesting that many FEF neurons are modulated by stimulus motion. Further, although FEF is known to *not* respond to low level visual features, in certain cases, similar to the task we analysed here, FEF neurons have been shown to be color-selective (Schall et al., 1995).

More precisely, when trained to utilize color in a target selection process, FEF neurons in monkeys were found to be selective to the color of the target (Bichot et al., 1996). These results are also in accord with studies that found sensitivity to motion direction in PFC in visual working memory tasks (Zaksas and Pasternak, 2006) and could also be related to our earlier finding that PFC color selectivity is weaker than motion selectivity (Buschman et al., 2012). Finally, although V4 is primarily known for shape and object processing (Roe et al., 2012), motion selectivity has also been observed in V4 neurons (Schmid et al., 2013).

Lastly, in LIP selectivity was not as clear as for the areas above. What was clear was that LIP was insensitive to the irrelevant categories in each task (deviations between brain and category DMs not significant). This underlines the importance of LIP in categorization. This is in accord with previous work by us and others. A recent study has suggested that LIP might be driving categorization in PFC (Swaminathan and Freedman, 2012) and our recent work also showed that coordinated interactions between LIP and PFC underlie categorization (Antzoulatos and Miller, 2016).

We then turned to computation selectivity. This is summarized in Figures 5 and 6. To understand this, we built deep RNNs. Although they comprised six LSTM layers (the same number of layers as the cortical network from which we recorded LFP responses), we use them only for *simulating brain computation, not* as precise descriptions of *anatomy*. We considered 2 variants of the same RNN. One trained to perform sensory processing and the other abstract categorization (sensory and category RNN respectively). We assumed that sensory processing would be based on low level visual features, while categorization would be based on information that the animal had learned after being trained to perform the task. Then we compared the RNN predictions to neural activity. We concluded that the computation a brain area performed would be similar to that of the RNN whose predictions were more similar to (had smallest deviations) and significantly correlated with brain activity. We trained them using LFPs as inputs and labels corresponding to different sensory stimuli or categories as outputs (depending on whether the RNN was processing sensory information or categorizing). Training RNNs with electrophysiological data (and to perform decision making tasks) is common, see e.g. (Rajan et al., 2016; Song et al., 2016).

RNNs in general and LSTMs in particular are appropriate for studying category representations in the brain because they are known to be able to learn temporal correlations very efficiently (Hochreiter and Schmidhuber, 1997; Pearlmutter, 1989)^10^. Temporal correlations in turn, have been suggested to link low level visual features into object categories in the context of binding theory (Milner, 1974; Von Der Malsburg, 1994). In earlier work, we have also hypothesized that synchronization might dynamically link category or stimulus selective neurons in decision making and memory tasks (Buschman and Miller, 2009; Buschman et al., 2012; Jones, 2016; Pinotsis et al., 2017).

, We did not consider task optimized networks. In future work, we will extend our results to such networks similar to the work by (Wang et al., 2018;Cadieu et al., 2014; Yamins et al., 2014).

To find its area’s preferred computation, we compared brain activity with the predictions of each of the two alternative RNNs (sensory and category). The fact that all six areas had been found to contain motion and color information in (Siegel et al., 2015) suggested that they could all be involved in performing motion and color processing and categorization^11^.

In the motion task, all areas (except PFC and FEF) preferred categorization. This suggests that computations in these areas focused on behavioural needs, to perform the two-choice motion categorization. For MT, this result is in accord with its well-known role during motion integration: MT neurons are able to recognise motion patterns by combining information from earlier visual areas (Rust et al., 2006). Also, several authors have suggested that hMT+/MT might be computing perceptual boundaries, similar to abstract categorization of motion direction that we found here (Bekhti et al., 2017; Hogendoorn and Verstraten, 2013; van Kemenade et al., 2014). In a recent study, (Bekhti et al., 2017) found that categorization occurred first in hMT+ and then this information was transferred to lower and higher areas. This can explain successful decoding of motion direction from early visual areas (Kamitani and Tong, 2006; Serences and Boynton, 2007). Also, this result fits with earlier results where LIP is known to integrate MT’s output up to a decision bound (Gold and Shadlen, 2007; Wimmer et al., 2015). Also, LIP could be sending signals back to feature-selective IT through feedback similar to that observed in selective attention studies (Squire et al., 2013).

FEF seemed to prefer sensory motion processing. This might be related to direction-specific eye-movement neurons that are abundant in FEF (Leigh and Zee, 2015). Here, they might encode the particular stimulus direction that the animal was attending to as part of some cognitive strategy to solve the task, possibly in conjunction with PFC (see below). PFC also seemed to prefer sensory processing. It might be receiving residual sensory motion input from FEF and other areas.PFC is known to process sensory motion signals as was also found by (Mante et al., 2013).

In the color task, all areas (except MT and PFC) were more involved in sensory color processing. This is in accord with our earlier work where we showed that processing in the color task is driven by sensory signals (Buschman et al., 2012). It also fits with several studies that have found successful color decoding using chromatic representations in early visual areas, e.g. (Brouwer and Heeger, 2009; Seymour et al., 2015). Similarly, it could be that computation of the perceptual boundary in the color task did not involve higher areas like the motion task. This could explain that evidence in favour of sensory processing was weaker in LIP (compared to earlier areas) and opposite in PFC.

In general, for both tasks, we found similar results using the two approaches considered here, domain and computation selectivity. This confirmed the validity of our analyses and suggests a way for understanding neural representations: compare brain responses to both a behavioural model (SDM/CDM) and deep neural network predictions and test if they give similar results. This can also guide the development of AI systems by designing deep neural networks whose layers perform computations similar to those performed by different brain areas that are involved in the same task.

All in all, representations changed flexibly depending on context (motion vs color task) and level of abstraction (sensory processing vs categorization). The motion task seemed to rely more on categorization, while the color task seemed to be driven by sensory computations. These results also fit well with several earlier findings in (Brincatt et al., 2018): (i) In that paper, motion and color information was found in ventral and dorsal stream areas and was included in distributed representations. If the brain uses this information to perform some computation, then this should be similar across many areas, which is what we found here. (ii) Also in that earlier paper, coding in most areas was found to reflect a mixture of sensory and categorical effects. Similarly, we found significant similarities between brain RDMs and RDMs from neural networks that perform both sensory processing and abstract categorization. (iii) Finally, according to (Brincat et al., 2018) categories arose gradually across the hierarchy. Our analysis, based on deep recurrent neural networks, revealed that gradual emergence is driven by sensory color and more abstract motion direction categorization.

Our results also fit well with earlier results by (Mante et al., 2013) who analysed dynamics from a similar perceptual discrimination task. That paper focused on PFC only and trained a single layer RNN on simulated input, not real LFP responses. Despite these differences, our results about PFC selectivity confirmed those earlier results. Like (Mante et al., 2013), we found that sensory information reaches PFC. Gating of sensory input is absent and filtering out of irrelevant (sensory) information by earlier brain areas did not occur. Also, (Mante et al., 2013) found that PFC responses during the motion and colour tasks occupy different parts of state space, and the corresponding trajectories are well separated along the axis of context (task). This can explain the flexible domain selectivity switching between tasks we found above. Recall that domain selectivity results from quantifying matches between brain responses and the geometry of stimulus and category domains. Differences in these matches require differences in brain responses. If responses did not occupy distinct parts of state space, then matches would not be different. We would not have observed domain selectivity. Based on this, we predict that besides PFC, responses of other brain areas should also occupy distinct parts of state space and their trajectories should be well separated.

All in all, our analysis sheds light to the biological basis of categorization and differences in selectivity and computations among different brain areas. It paves the way for constructing neural networks that can replicate brain dynamics underlying complex sensorimotor decision making tasks. Elucidating such differences can be important for building automated systems for intelligent decision making in multidimensional domains, like driverless cars, pilot support systems, medical diagnosis algorithms etc. We hope our work can help make progress in this direction.

## Acknowledgements

This work was supported by NIMH R37MH087027. DAP acknowledges illuminating discussions and useful comments by Dr Pouya Bashivan.

## Footnotes

1 Trials with shorter RT than the average could include visual and/or premotor responses as confounds. Automated decision making systems would need to decide based on such confounds. Thus we also analyzed data with the same confounds. We chose not to exclude them, e.g. not to model activity until the RT specific to each trial, because that would be irrelevant for automated systems.

2 Deviations can be intuitively thought of as distances in a multidimensional space of second order differences. This is the reason for choosing to compute (*1-correlation*) as opposed to *correlation* as discussed in (Kriegeskorte et al., 2008).

3 We used N=100 repetitions as this was the number used in Kriegeskorte et al., 2008. The value of N does not change our conclusions which also employ post hoc analyses based on multilinear regression.

4 Because we analyzed activity until average RT that could contain visual and/or premotor responses (see footnote 2), task dependent FEF selectivity could be due to *saccade selective*, not motion or color selective, brain activity.

5 MT and V4 selectivity appear stronger during the color task (smaller deviations in color vs motion task). However, because we are using different neural data (trials) in each task, we cannot estimate relative selectivity strengths in the two tasks. Smaller deviations could be attributed to other confounds present in the two datasets and *not* to genuine brain area selectivity.

6 We trained the network separately for the motion and color tasks. Our focus was on computations performed by each area during each task independently, not switching tasks *per se*. Also, separate network training results in correlations (between brain activity and network predictions) that are not biased in favor of one domain (either motion or color). Had we trained the network on both domains, RDM analysis below could have favored the domain for which discrimination by the brain was relatively better. Focusing on each domain independently allowed us to tease out differences between sensory processing and categorization in that domain; as opposed to conflating differences in the level of abstraction with differences in perception of these two domains by the brain.

7 Instead of 2 SDMs and 2 CDMs we here have 6 network RDMs, one for each layer. Thus, we have 6×2 bars (for the 2 deep RNN variants performing sensory processing and categorization) instead of 4 bars, see next paragraph.

8 Besides the input layer because this does not predict brain activity.

9 Note *ΔBICsc>6* vs *ΔBICsc>10* for the color and motion tasks respectively. Thus, relative evidence in favor of categorization is stronger in the motion task. Given this result and the earlier result (Figure 4) that MT was selective to the motion domain, it seems MT prefers more motion than color categorization. However, we cannot exclude that relative differences in computation selectivity in the two tasks are due to confounds, not intrinsic MT selectivity.

10 In Supplementary Results, we also show that LSTMs predicted neural responses better than linear regression.

11 Whether all brain areas are necessary for task computations is beyond the scope of our paper. This would require a separate analysis, like a lesion study.

